# Science-wide mapping of institutions based on affiliated authors’ impact and research integrity proxies

**DOI:** 10.64898/2025.12.16.694665

**Authors:** John P.A. Ioannidis, Jeroen Baas, Roy Boverhof, Cyril Voyant

## Abstract

We aimed to generate institution-level data of indicators that may reflect, as proxies, the impact and the research integrity of the authors affiliated with them and that also adjust for institution size and scientific subfields where these authors work. We used the Scopus database to generate institution-level data on top-cited authors (according to a composite citation indicator) and three indicators that may reflect problems with research integrity: the total number of authorships in papers retracted for reasons other than publisher/journal errors; and subfield-adjusted proportions of top-cited authors who have high (>95% percentile) self-citation rates or high (>95% percentile) rates of publications in Scopus-discontinued titles. There was modest overlap among top institutions based on volume and based on number of top-cited authors. However, there was minimal overlap when institutions with the highest proportions of top-cited authors were considered; these included mostly research institutes and tech organizations and only a minority of universities. Institutions with the highest self-citation, discontinuation, and retraction penalties clustered in specific countries such as Saudi Arabia, China, Malaysia, Iran, India and Indonesia. The publicly available datasets that we provide allow the exploration of institution-level research assessments that balance high impact against proxies of diverse aspects of research integrity.

## INTRODUCTION

Evaluations of institutions are widely popular and result in diverse appraisals and rankings that often attract justifiable criticism as to their reliability and face validity (1–4). Most evaluations typically try to measure impact using different indicators of excellence, influence, reputation, and recognition. In most of these assessments, there is very limited or no attention to issues of research integrity. For example, rankings that focus on impact typically do not account at all for misconduct that may lead to retractions. They are also typically entirely silent to spurious practices that may lead some scientists to high publication and citation counts through gaming. Another fundamental problem with such assessments is that information on how many people are employed by a university or other institution is difficult to collect in a standardized way across institutions worldwide. There are many different types of faculty and staff, e.g. considering tenured and non-tenured, different ranks, regular and adjunct appointments, let alone hierarchies in universities versus other organizations that publish research. The exact categories and which ones are counted by each institution may vary a lot. This heterogeneity generates biases when comparing different institutions directly and when normalizing impact metrics by the number of affiliated researchers.

Several assessment systems try to capture the number of people affiliated with each institution that have achieved some specific hallmark of impact, e.g. Nobel prizes, or other major recognitions. However, for the most globally acknowledged – and thus by definition rare – recognitions only few institutions have any such individuals. Many assessment systems end up using numbers of published papers, citations, or numbers and proportion of papers in percentiles of citations. Absolute numbers of papers and citations, however, would again require some adjustment for the number of available producing authors to be more meaningfully interpretable. Proportion of papers in citation percentiles does achieve some reasonable adjustment (5) but provides no information on institutional size and on whether the high-impact work is produced by a few individuals or spread across many. Moreover, diverse research integrity issues remain completely ignored and unaccounted for.

Here, we use the Scopus database (6) and previously derived datasets of scientific subfield-adjusted lists of top-cited scientists (7–10) to derive standardized, institution-level metrics that may be used for appraising the volume and impact of diverse institutions across science and around the world. For each institutional affiliation, we leverage these resources to generate information on the overall number of publishing authors and the number of publishing authors who fulfill pre-defined threshold criteria for onset of their publication age, their overall publication volume, and their overall volume of publications as single, first, or last authors. Instead of selecting institution-specific categories of employees, we count all authors with an institutional affiliation, and separately specifically those who have started publishing in the last 45 years (a typical duration of a scientific career) and who have demonstrated substantial productivity. Moreover, for each institution we obtain the number of authors who are top-cited in their scientific subfield overall and after using the same filtering thresholds. The proportion of top-cited authors among all authors with such filtering thresholds may then be considered as a first adjusted benchmark of impact for each institution. Finally, we propose that impact should not be examined alone but seen in combination with indicators that may be proxies of diverse aspects of research integrity. Therefore, we add further adjustments that consider for each institution the proportion of its eligible top-cited authors who have high self-citations and high share of their papers published in journal titles that have been discontinued for quality concerns by Scopus; and, furthermore, the total authorships of retracted papers not due to publisher/journal error. These adjustments aim to balance the citation impact against indicators that may reflect, as proxies, diverse aspects of research integrity. Strengths and caveats of such an approach to assessing institutions are then discussed and put into perspective.

## METHODS

### Scopus database

We used the Scopus database (6) with a data freeze on August 1, 2025 (11). Similar to previous work (7–10), we limited all analyses to authors who have published at least 5 full papers in the database (defined as articles, reviews, and conference papers in Scopus), yielding a total of 10,933,183 eligible authors. The accuracy of Scopus for assigning all papers to the correct author and not splitting papers of the same author to different author files is deemed very high. The process is reviewed in (6). A study found precision and recall of 99% and 98%, respectively for Japanese researchers (12), while an average precision of 98.1% and an average recall of 94.4% was reported in a 2020 internal overview of the database (6).

### Aggregation of organizations

The merging of author affiliations to “aggregated” organizations (which we refer to as *institutions*) is done according to the Scopus process for assigning one primary affiliation to each author and for creating hierarchies of organizations that are mapped to each specific institution. Details are discussed in Appendix Methods. The full hierarchy of organizations mapped to each institution can be found in the subscription edition of Scopus.

### Top-cited authors

For each organization and institution, the number of top-cited authors is counted according to the latest edition (August 2025) of the citation indicators databases (11) that include all authors who are in the top-2% of their scientific subfield (across 174 subfields classified by Science-Matrix (13) or among the top 100,000 of all science based on a composite citation indicator. The composite citation indicator combines 6 citation indicators, including total citations, Hirsch h-index, Schreiber co-authorship-adjusted hm-index, citations to single-authored papers, citations to single- or first-authored papers, and citations to single-, first- or last-authored papers (7–10). It aims to accommodate co-authorship and indirect measures of author contributions through author positions since reliable, detailed, standardized data on author contributions are not available across the scientific literature. Some fields use alphabetic listing, in which case first and last author positions do not carry particular importance; however, we have previously estimated that alphabetic listing may affect only a small minority of papers (7). Authors are considered in the top-cited list if they manage to pass the selection thresholds in analyses excluding self-citations or in analyses including self-citations. Moreover, separate datasets of top-cited authors are generated for career-long citation impact and for citation impact limited to the citations received in the single most recent year (here, this is calendar year 2024). Of the 10,933,183 authors with at least 5 full papers in Scopus, a total of 230,333 and 236,313 are captured as top-cited in the career-long and single recent-year datasets, respectively. In the Results, we first focus on career-long data, since they reflect cumulative long-term influence. The single recent-year analyses may be more vulnerable to gaming, e.g. through fake papers and orchestrated self-citation cartels, and they are presented separately and briefly compared with career-long results.

### Threshold filters for authors and proportions of top-cited authors

We have derived data for the number of authors and the respective number of those who are included in the datasets of top-cited authors using threshold filters for publication age and for productivity volume and patterns.

For publication age, we used a filter for starting to publish (any item that is Scopus-indexed) in 1980 or later. This allows for up to 45 years of publication history, excluding authors who started publishing in earlier years. Assuming one publishes their first paper usually in their 20s to early-30s, a 45-year window covers the time to typical retirement for a full academic or other research career. We also derived data for sensitivity analyses where we further excluded authors who have not had any Scopus-indexed items published from 2020 onwards. These sensitivity analyses further exclude authors who may be dead or may be scientifically inactive.

For productivity threshold filters, we used one filter based on the number of total Scopus-indexed published items and another one for single-, first-, or last-authored items. For the number of total published items, we derived data for the number of authors with 40 or more total published items. Authors with <40 published items have negligible chances of being inducted in the top-cited list (97% of the top-cited authors have 40 or more published items). For the filter of single-, first- or last-authored items, we set the filter at having 5 or more such publications; 99.7% of the top-cited authors have 5 or more such publications.

We derived the proportions of authors who are included in the datasets of top-cited authors for career-long impact and separately for single recent-year impact. When all authors are considered without further filtering for publication age, many included authors may represent individuals that reflect the long past legacy of the institution; this may affect to a different extent institutions with older versus only recent history. When no filter for the number of total published items is set, institutions who are more actively engaging many early career and in-training researchers may have a large denominator of total eligible authors than those that have few such researchers and a larger share of well established, senior scientists. Early career and in-training researchers have minimal chance of reaching top-cited status so fast (14), therefore the proportion of top-cited authors would penalize institutions with many such researchers unless they are filtered out in the calculations. Finally, when no filter is set for a minimum number of single-, first- or last-authored papers, authors who have not much of a trajectory of leading roles in research may also distort the percentage of top-cited authors, since these authors rarely make it to the lists of top-cited authors based on the composite citation indicator used. As above, the composite citation indicator placed major emphasis on single-, first- or last-authored contributions.

In the Results section, we present first data on all authors with at least 5 Scopus-indexed full papers and then we focus primarily on the subset of those who started publishing in 1980 or later, have published 40 or more Scopus-indexed items in their career, and these include at least 5 single-, first- or last-authored items. The selected subset for this analysis (henceforth called *primary analysis*), may approximate the group of senior active faculty in universities or equivalent leading roles in other institutions and may be most appropriate for standardized institutional comparisons rather than the all-inclusive authors’ data.

### Analyzed institutions

A very large number of organizations may have very few authors assigned to them even at the aggregated institutional level. Any evaluation of the number or proportion of highly cited authors in them would not be meaningful given the sparse information. Therefore, we first generated data focused on only those institutions that included at least 50 authors who have had at least 15 Scopus-indexed items. These data are made available in public repositories (http://doi.org/10.17632/jn5j7gjpzj.1) for further exploration by interested users. However, we realized that this filter includes many institutions with too few authors meeting the primary analysis criteria. Therefore, we finally generated percentile rankings focused exclusively on institutions that have at least 100 authors eligible by the primary analysis filters.

### Indicators that may be proxies of diverse aspects of research integrity

The research integrity of authors and their institutions can be evaluated with diverse indicators that capture different, potentially complementary aspects. These indicators function as proxies rather than firm proof of research misconduct or gaming practices. Each of them may offer some insights into research practice and malpractice and consideration of multiple indicators may offer a more comprehensive view rather than focusing on a single one. Here we used three indicators that capture features that are previously recognized to reflect directly or indirectly problems in research integrity and where there is access to science-wide data: exceedingly high rates of self-citations, publications in titles that have been discontinued due to quality concerns and retractions due to author issues (rather than publisher/journal error).. Illustratively, recently they were incorporated on a system of institutional integrity ranking, the Research Integrity Risk Index (15). Moreover, science-wide data is available for these indicators, as described below.

We have previously linked retraction data for each author in the Retraction Watch database (16) with the Scopus-based author citation datasets with (17,18). As described previously, we consider only retractions not due to publisher/journal errors and these represent the majority of catalogued retractions (16–18). Top-cited authors are few for most institutions and authors with retractions may not reach or may fall off the range of being top-cited. Therefore, we consider all institutional authors who started publishing in 1980 or later and had at least 5 published items, we count for each the number of their publications not due to publisher/journal error and then sum all these retraction authorships, ΣR. Illustratively, a single retracted paper with three institutional authors contributes 3 retraction authorships for that institution.

Self-citations may often be fully appropriate to put new work in the context of older work. Moreover, authors in earlier career stages may appropriately have higher self-citation rates. Therefore, we focused only on top-cited authors, Since it is impossible to evaluate in depth whether each self-citation is appropriate, for each top-cited author we note whether their percentage of self-citations is higher than the threshold of the 95% percentile for self-citations in the respective subfield among the top-cited author peers. We already generate such data on self-citations and subfield thresholds (considering self-citations from all authors of a given paper) in the annually updated citation datasets (and high rates of self-citations (10,19) and here we used the August 2025 edition (11).

In a similar fashion, for each top-cited author we note whether their percentage of published items that appeared in titles that have been discontinued by Scopus is higher than the 95% percentile for items in discontinued titles among the top-cited author peers in the respective subfield. Discontinuation of a title here means that after a title is selected for re-evaluation and flagged for discontinuation by the Content Selection and Advisory Board (CSAB) (6), new content will no longer be included in Scopus. Discontinuation means that the title no longer meets the quality standards set by CSAB. For details on how quality is appraised see (20). Already indexed content from this title will remain in Scopus unless severe unethical publication practices can be proven, in which case indexed content may be removed (a very rare occurrence). Journals that simply cease to publish are not considered “discontinued”; the term is reserved for active decisions by the CSAB.

### Summary score considering both impact and research integrity proxy indicators

For each institution, we generate a score that combines information on both the citation impact and research integrity proxy indicators. This score is given by

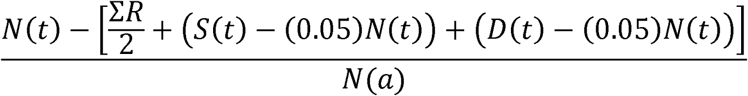

where N(t) is the number of top-cited authors in the primary analysis, N(a) the number of all authors in the primary analysis, S(t) the number of top-cited authors in the primary analysis with self-citations above the 95% percentile, D(t) the number of top-cited authors in the primary analysis with percentage of items published in discontinued titles exceeding the 95% percentile, and ΣR the sum of all retraction authorships across all institutional authors with at least 5 published items who started publishing in 1980 or later.

In this framework, retractions by any author always penalize an institution and the loss due to having two retracted authorships is equivalent to the gain from having a top-cited author. Conversely, the proportion of its top-cited authors with high self-citations and high rates of publications in discontinued titles may offer a bonus or penalty to the institution depending on whether they are less or more than 5%.

Of note, retraction authorships are counted for all eligible authors in an institution, while the other two proxy indicators focus on the eligible top-cited authors only. The reason for this choice is that authors with retracted work not due to publisher/journal error may suffer a considerable citation penalty (21,22) that keeps them away or removes them from the top-cited list, while there is no evidence for such a citation penalty for those with high values on the other two proxy indicators.

### Percentile ranking of institutions

These summary scores were used to generate institutional percentile rankings. We also performed extensive sensitivity analyses of the institutional percentile ranking, by varying the main tuning parameters of the metric. Specifically, we changed the self-citation and discontinued-titles threshold percentiles (p = 80, 95, 99) and divided the contribution of retraction authorships ΣR by factors f = 1, 2 (baseline), and 4. For each combination of (p, f), we recomputed the adjusted score for institutional percentile ranking using

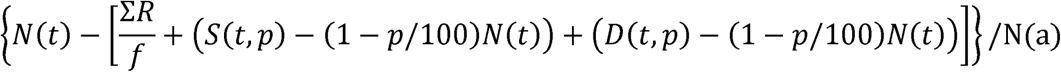

where N(t) is the number of top-cited authors in the primary analysis, ΣR is the number of retraction authorships among eligible authors, S(t,p) is the number of top-cited authors whose self-citation percentage exceeds the p-th percentile in their subfield, and D(t,p) is the number whose share of publications in discontinued titles exceeds the subfield-specific p-th percentile. We then quantified agreement with the baseline (p = 95, f = 2) ranking using Spearman rank correlations for both the institutional percentile ranks and the adjusted percentage of top-cited authors. To visualize potential shifts, we generated scatter plots and Bland-Altman plots of the baseline versus sensitivity-analysis rankings and adjusted percentages. We also estimated smoothed distributions using kernel density estimation (KDE) to visualize the distribution of adjusted institutional scores across specifications for each choice of (p, f).

## RESULTS

### Overall data with institutions

Data are provided in the publicly accessible link (http://doi.org/10.17632/jn5j7gjpzj.1) on a total of 6,979 institutions with ≥50 authors who have ≥15 Scopus-indexed items. These institutions cover 78% of all authors with at least 5 full publications (8,524,471/10,933,183) and the vast majority of career-long top-cited authors (90.0% [207,256/230,333]). When limited to authors eligible for the primary analysis (those who started publishing in 1980 or later, have published 40 or more Scopus-indexed items in their career, and these include at least 5 single, first- or last-authored items), there were 1,214,702 authors across these 6,979 institutions, of which 145,704 (12%) were top-cited according to career-long impact. 22,823 authors (2,781 top-cited ones) were counted for more than one aggregated affiliation.

Of the 6,979 institutions, 2,380 have also at least 100 authors who started publishing in 1980 or later, have published 40 or more Scopus-indexed items in their career, and these include at least 5 single-, first- or last-authored items and are thus used for presented percentile rankings.

### Institutions with highest numbers of top-cited authors

Table 1 and Supplementary Table 1 present information on the 100 institutions that have the highest number of top-cited authors in the primary analysis (range 278-1,135 top-cited authors per institution). The listed institutions are highly recognizable organizations with undisputable major presence in science. However, they vary a lot on the proportion of top-cited authors in the primary analysis from 6.98% (Shanghai Jiao Tong University) to 40.13% (University of California, Berkeley). Supplementary Table 1 also shows, for comparison, data for these institutions considering all authors. There is substantial variation in the share of authors eligible for the primary analysis (range 9-26%) with the lowest values at UC Berkeley and MIT. Variation is even greater in the proportion of top-cited authors who meet the primary-analysis criteria (range 46-99%). Chinese institutions and to a lesser degree also institutions in Singapore and South Korea have almost all their top-cited authors qualify for the primary analysis. This is because in contrast to American, European, and Australian institutions, Asian institutions have seen massive growth more recently and thus very few of their top-cited authors are excluded because of old (pre-1980) start of publication history. Nevertheless, Chinese institutions still display the lowest proportion of top-cited scientists within Table 1 and Supplementary Table 1.

**Table 1.** Number of authors and of top-cited authors (career-long citation impact) in the 100 institutions with the highest number of top-cited authors in the primary analysis. Data per region, with details shown on top-20 institutions. Detailed data for all 100 institutions appear in Supplementary Table 1.

| Region | Number among top 100 | Among top 20 |
| --- | --- | --- |
| United States | 47 | Harvard University (1135), University of Michigan, Ann Arbor (913), Stanford University (912), Johns Hopkins University (782), University of Pennsylvania (781), University of Washington (751), University of California, Los Angeles (687), Columbia University (633), University of California, San Diego (629), Mayo Clinic (592), Duke University (590), Massachusetts Institute of Technology (564), University of Minnesota Twin Cities (551), Cornell University (544) |
| United Kingdom | 12 | University of Oxford (828), University College London (810), University of Cambridge (651), Imperial College London (542) |
| France | 7 | None |
| China | 5 | None |
| Australia | 5 | None |
| Canada | 4 | University of Toronto (1038), The University of British Columbia (566) |
| Japan | 3 | None |
| Netherlands | 3 | None |
| Denmark | 2 | None |
| Belgium | 2 | None |
| Switzerland | 2 | None |
| Germany | 2 | None |
| Sweden | 2 | None |
| Singapore | 1 | None |
| Korea, Republic of | 1 | None |
| Hong Kong | 1 | None |
| Finland | 1 | None |

### Institutions with highest numbers of authors

Table 2 and supplementary Table 2 present information on the 116 institutions that have more than 10,000 authors (and up to 35,547) with 5 or more full publications. These institutions have between 606 and5,027 authors eligible for the primary analysis, with the lowest value observed at Pfizer, Inc. – apparently few authors with pharma affiliation publish many papers. The number of top-cited authors varies vastly, from 14 (Southern Medical University) to 1894 (Harvard University) for all top-cited authors and from 14 to 1135 in the primary analysis. The proportion of top-cited authors in the primary analysis also showed wide variation from 1.05% (Capital Medical University) to 40.13% (UC Berkeley). Several institutions in this list of the largest institutions have very low proportions of top-cited scientists. In 13 institutions, this proportion is below 6% in the primary analysis, including 8 institutions in China and others in Russia, India, Mexico, Czech Republic, and Brazil.

**Table 2.** Number of authors and of top-cited authors (career-long citation impact) in the 116 institutions with over 10,000 authors. Data per region, with details shown on top-20 institutions. Detailed data for all 116 institutions appear in Supplementary Table 2.

| Region | Among the 116 | Among top 20 |
| --- | --- | --- |
| United States | 38 | Harvard University (26366), Stanford University (24041), University of Michigan, Ann Arbor (22335), Johns Hopkins University (21268), University of Pennsylvania (20953), University of California, Los Angeles (20358), University of Washington (19683) |
| China | 27 | Shanghai Jiao Tong University (35547), Zhejiang University (31288), Peking University (27753), Tsinghua University (24487), Huazhong University of Science and Technology (24022), University of Chinese Academy of Sciences (23373), Sichuan University (23245), Fudan University (21490), Central South University (20139), Jilin University (18882) |
| France | 8 | None |
| United Kingdom | 7 | None |
| Germany | 4 | None |
| Japan | 4 | The University of Tokyo (18748) |
| Canada | 4 | University of Toronto (30477) |
| Switzerland | 2 | None |
| Sweden | 2 | None |
| Belgium | 2 | None |
| Italy | 2 | None |
| Australia | 2 | None |
| Korea, Republic of | 2 | None |
| Netherlands | 2 | None |
| Taiwan, Province of China | 1 | None |
| Finland | 1 | None |
| Norway | 1 | None |
| Czechia | 1 | None |
| Russian Federation | 1 | None |
| Singapore | 1 | None |
| India | 1 | None |
| Denmark | 1 | None |
| Brazil | 1 | Universidade de São Paulo (23723) |
| Mexico | 1 | None |

### Institutions with highest percentage of top-cited authors

Table 3 and Supplementary Table 3 show the 100 institutions with the highest proportions of top-cited authors (range, 30.43-79.17%) among all 6,979 institutions. The top-20 of Tables 1 and 3 have no institutions in common. Most of the Table 3 institutions are small- or modest-size organizations, and many of them are research institutes and companies in the tech space. The top-100 include only 30 universities and polytechnics (Supplementary Table 3).

**Table 3.** Number of authors and top-cited authors (career-long impact) in institutions with the highest percentage of top-cited authors in the primary analysis. Data per region, with details shown on top-20 institutions. Detailed data for all 100 institutions appear in Supplementary Table 3.

| **Region** | **Number among top 100** | **Among top 20** |
| --- | --- | --- |
| United States | 38 | Meta FAIR (69.23%), Institute for Advanced Study (64.71%), National Bureau of Economic Research (57.14%), California Institute for Quantitative Biosciences (50.00%), Santa Fe Institute (50.00%), Howard Hughes Medical Institute (42.86%), Princeton University (41.93%), National Center for Health Statistics (41.86%) |
| Germany | 13 | Max-Planck-Institut für Eisenforschung GmbH (47.62%), CISPA - Helmholtz Center for Information Security (46.15%), Max Planck Institute for Coal Research (43.33%), Max Planck Institute for Informatics (42.11%) |
| United Kingdom | 13 | The Medical Research Council Laboratory of Molecular Biology (47.17%) |
| Spain | 8 | CSIC - Instituto de Ciencia y Tecnología de Alimentos y Nutrición (ICTAN) |
|  |  | (40.38%) |
| Netherlands | 3 | Wolters Kluwer (79.17%) |
| Canada | 3 | Perimeter Institute for Theoretical Physics (58.62%) |
| Taiwan,<br>Province of<br>China | 2 | None |
| France | 2 | INSEAD, Europe (56.25%), EURECOM (42.42%) |
| Switzerland | 2 | None |
| Italy | 2 | None |
| Hong Kong | 2 | None |
| Türkiye | 2 | None |
| Ukraine | 1 | None |
| China | 1 | None |
| Kuwait | 1 | None |
| Palestine,<br>State of | 1 | None |
| Norway | 1 | None |
| Iraq | 1 | None |
| Iran, Islamic<br>Republic of | 1 | None |
| India | 1 | National Aerospace Laboratories India (42.11%) |
| United Arab<br>Emirates | 1 | Mohamed Bin Zayed University of Artificial Intelligence (44.00%) |
| Austria | 1 | None |

For institutions with a small number of authors in the primary analysis, their prominently high placement may be spurious or occasionally may even point to gaming practices, since a few gamed top-cited authors can propel a small institution to high placement. To avoid challenges with small numbers of eligible authors, Table 4 and Supplementary Table 4 show the top-100 institutions with the highest proportion (24.84-41.93%) of top-cited authors in the primary analysis, when limited to the 2,380 institutions with at least 100 authors eligible for the primary analysis. The top-20 in Tables 1 and 4 have little overlap.

**Table 4.** Number of authors and top-cited authors (career-long impact) in institutions with the highest percentage of top-cited authors in the primary analysis limited among the institutions with at least 100 authors for the primary analysis. Data per region, with details shown on top-20 institutions. Detailed data for all 100 institutions appear in Supplementary Table 4.

| Region | Number among top 100 | Among top 20 |
| --- | --- | --- |
| United States | 62 | Princeton University (41.93%), University of California, Berkeley (40.13%), Carnegie Mellon University (38.22%), Alphabet Inc. (37.29%), University of California, Santa Barbara (36.38%), Massachusetts Institute of Technology (34.16%), Rockefeller University (31.93%), Georgia Institute of Technology (31.77%), Rice University (31.36%), University of Maryland, College Park (30.91%), University of Illinois Urbana-Champaign (30.22%), Microsoft Corporation (30.20%), The University of Texas at Austin (30.11%) |
| United Kingdom | 17 | The James Hutton Institute (33.98%), London School of Economics and Political Science (32.41%), University of Cambridge (31.86%), University of Oxford (30.66%) |
| Hong Kong | 5 | Hong Kong University of Science and Technology (31.83%), The Education University of Hong Kong (30.53%) |
| Canada | 3 | None |
| Israel | 3 | None |
| Switzerland | 2 | ETH Zürich (32.37%) |
| China | 1 | None |
| Netherlands | 1 | None |
| Türkiye | 1 | None |
| Ireland | 1 | None |
| Spain | 1 | None |
| Singapore | 1 | None |
| Germany | 1 | None |
| Saudi Arabia | 1 | None |

### Retractions, self-citations, and discontinued titles

The publicly deposited files (http://doi.org/10.17632/jn5j7gjpzj.1) provide information for each institution on retraction authorships ΣR across all authors who had at least 5 full published papers and started publishing in 1980 or later, top-cited authors in the primary analysis with high self-citations S(t,p), and top-cited authors in the primary analysis with high rates of items published in discontinued titles D(t,p). Many institutions have high values when contrasted to the number of top-cited authors and to all eligible authors in the primary analysis, suggesting that their ranking will be markedly affected when these parameters are also considered.

Prevalence of institutions among the adjustment components shows large differences by country. When we divide each component-specific penalty by the number of top-cited primary analysis authors N(t) for each institution, we get the relative impact of the adjustments of each component on the institution. For retractions, the top 100 institutions ranked by retraction adjustment per N(t) are heavily concentrated in China (75/100 institutions). These 75 China institutions with high retractions account for 18% of all Chinese institutions among the 2,380. Other countries with at least 2 institutions in the top 100 of retractions were Indonesia (n=3, 19% of its total institutions), Malaysia (n=2, 10%), Morocco (n=2, 25%) and South Korea (n=2, 3%). For self-citation, the top 100 institutions ranked by the self-citation adjustment per N(t) are dominated by Russia (n=26, 52% of its total institutions), China (n=9, 2%), Indonesia (n=7, 44%), Poland (n=6, 15%) and the Czech Republic (n=6, 35%). For discontinued titles, the top 100 institutions ranked by the discontinued-title adjustment per N(t) are led by India (n=12, 13%), China (n=12, 3%), Malaysia (n=9, 45%), Saudi Arabia (n=6, 32%) and Russia (n=4, 8%).

### Percentile ranking after adjustment for research integrity indicators

Table 5 and Supplementary Table 5 show the top-100 institutions according to the percentile ranking that corrects for retractions, self-citations and discontinuations when limited to the 2,380 institutions with at least 100 authors who started publishing in 1980 or later, have published 40 or more Scopus-indexed items in their career, and these include at least 5 single, first-,or last-authored items. For the 2,380 institutions, their percentile ranking is available in the publicly deposited files (http://doi.org/10.17632/jn5j7gjpzj.1). As shown, the adjusted percentile ranking for many institutions is markedly different from the unadjusted percentile ranking using only the proportion of top-cited authors.

**Table 5.** Number of authors and top-cited authors (career-long impact) in institutions with the highest percentage of top-cited authors after correction, in the primary analysis limited among the institutions with at least 100 authors for the primary analysis. Data per region, with details shown on top-20 institutions. Detailed data for all 100 institutions appear in Supplementary Table 5.

| Region | Number among top 100 | Among top 20 |
| --- | --- | --- |
| United States | 62 | Alphabet Inc. (40.20%), Carnegie Mellon University (37.73%), University of California, Berkeley (37.26%), Princeton University (36.47%), Massachusetts Institute of Technology (32.97%), Microsoft Corporation (32.61%), University of California, Santa Barbara (32.24%), Georgia Institute of Technology (31.35%), Woods Hole Oceanographic Institution (31.03%), National Institute of Diabetes and Digestive and Kidney Diseases (NIDDK) (30.11%), Rockefeller University (30.08%), Rice University (29.74%), Colorado School of Mines (28.65%), University of Illinois Urbana-Champaign (27.59%) |
| United Kingdom | 15 | The James Hutton Institute (36.41%), University of Cambridge (28.10%), University of Oxford (27.61%) |
| Canada | 4 | None |
| Hong Kong | 4 | Hong Kong University of Science and Technology (31.08%) |
| Switzerland | 2 | ETH Zürich (31.05%), École Polytechnique Fédérale de Lausanne (29.19%) |
| Israel | 2 | None |
| Netherlands | 2 | None |
| Ireland | 1 | None |
| Türkiye | 1 | None |
| France | 1 | None |
| Italy | 1 | None |
| Spain | 1 | None |
| Japan | 1 | None |
| Sweden | 1 | None |
| Denmark | 1 | None |
| Germany | 1 | None |

As shown in Table 6, China has the largest number of institutions (n=410) among the 2380 compared with any other country, followed by the USA (n=344), while Japan, France, and United Kingdom also have more than 100 institutions each in that list. A total of 28 countries/regions have at least 15 institutions among the 2380. United Kingdom, Netherlands, Switzerland, Canada, United States, Australia and Sweden have very high percentile rankings of their institutions with very similar median percentile values (82.6th-85.5th percentile). Conversely Saudi Arabia, China, Malaysia, Iran, India and Indonesia have very low percentile rankings of their institutions, with median values ranging from 3.9th to 21st percentile. These countries are penalized the most from the three adjustments, with the median cumulative adjustment exceeding their number of top-cited authors N(t) by 1.29-2.75-fold.

**Table 6.** Regions with at least 15 institutions among the 2,380 institutions with >=100 primary-analysis authors: counts, median adjusted percentile, and median cumulative adjustment (as proportion of N(t)). Detailed data for all 2,380 institutions appear in the supplementary excel file sheet Raw Data.

| Region | Number of institutions among the 2,380 | Median percentile ranking among the 2,380 (using adjusted baseline) | Median cumulative adjustment (as proportion of $N(t)$) |
| --- | --- | --- | --- |
| China | 410 | 14.3 | 2.06 |
| United States | 344 | 83.6 | 0.13 |
| Japan | 152 | 43.0 | 0.39 |
| France | 137 | 58.5 | 0.18 |
| United Kingdom | 117 | 85.5 | 0.10 |
| Italy | 96 | 50.8 | 0.34 |
| Germany | 95 | 74.8 | 0.18 |
| India | 92 | 18.8 | 1.29 |
| Spain | 80 | 50.1 | 0.21 |
| Korea, Republic of | 71 | 36.7 | 0.62 |
| Russian Federation | 50 | 27.9 | 0.82 |
| Brazil | 47 | 33.5 | 0.53 |
| Australia | 46 | 83.2 | 0.13 |
| Türkiye | 46 | 33.1 | 0.73 |
| Canada | 42 | 84.0 | 0.09 |
| Poland | 41 | 43.0 | 0.38 |
| Iran, Islamic Republic of | 34 | 15.4 | 1.70 |
| Taiwan, Province of China | 32 | 56.6 | 0.40 |
| Malaysia | 20 | 15.3 | 1.61 |
| Saudi Arabia | 19 | 3.9 | 2.75 |
| Netherlands | 18 | 84.5 | 0.08 |
| Switzerland | 17 | 84.5 | 0.14 |
| Czechia | 17 | 38.3 | 0.60 |
| Romania | 17 | 26.6 | 0.90 |
| Finland | 16 | 73.8 | 0.09 |
| Indonesia | 16 | 21.0 | 2.01 |
| Sweden | 15 | 82.6 | 0.12 |
| Austria | 15 | 71.5 | 0.22 |

### Single recent-year analyses and sensitivity analyses

Analyses using the single recent-year data and sensitivity analyses are presented in Appendix Results.

## DISCUSSION

We provide data that allow standardized assessment of institutions worldwide based on the number of publishing authors with each specific institutional affiliation; the concentration of top-cited authors in each institution, and three indicators that may offer proxies of diverse research integrity aspects (retractions and high self-citation and publication at discontinued title rates).

The percentage of top-cited authors can be calculated for different sets of authors. Here, we preferred for the primary analysis a set that considers only authors whose publication activity spans a typical research-career window and who have a substantial publication record and at least 5 published items with leading and/or senior authorship positions. Such data allows the use of a standardized process for calibrating institutional impact through the proportion of its top-cited authors. Defining the set of authors based on objective publication age and productivity criteria regardless of what their academic or other rank and status creates some common ground for comparisons of very diverse institutions. Academic positions, ranks, and status would be extremely difficult to standardize across thousands of diverse institutions. These institutions include not only universities (that already have different systems for classifying faculty and staff) but also many other organizations and entities that publish scientific papers.

Science-wide mapping of nearly 7,000 institutions reveals that top-cited scientists are disproportionately concentrated in high numbers in a few institutions. However, there are many other institutions of more modest size or even relatively small size that have high proportions of top-cited authors among their ranks. Without adjusting by size, the excellent profiles of these institutions would have been buried under the volume of the largest ones and remained unnoticed. Institutions with high values include not only large universities, but also research institutes of more modest size, and several tech companies and other organizations. Conversely, among the very largest institutions (with size determined by their numbers of total authors), there is extremely large variability as to the presence and percentage of top-cited scientists.

Our databases may be used for research assessments of institutions. However, we caution that, in principle, institutional rankings may be problematic as extensively discussed before (1–4, 23–25). Frustrated with the misuse of institutional rankings, some initiatives such as COARA have called even for their total avoidance (4). Instead of total abolition a more pragmatic operationalization is to use ranking metrics more judiciously (26), generate more standardized metrics (27), and consider also systematic appraisals of gaming processes (28,29) to better accommodate research integrity. The three proxy indicators of research integrity may best be seen as offering warning signals rather than validated proof of research integrity failure at an institutional level. The same cautious principles apply also to interpreting the summary score ranking which should be seen as heuristic rather than some gold standard of validated ranking. If our data are used for any assessment and ranking purposes, they should be strictly interpreted for what they show. Clearly, they may not capture sufficiently many other dimensions that are important for understanding the impact of research and other organizations, e.g. education, policy, community work, health services, and more.

Our analysis also reveals that focus on top-cited authors may be one-sided and may not account for aspects that touch on research integrity. We demonstrate that some institutions (mostly small ones) from countries with very limited resources have extremely high percentages of top-cited authors. This pattern should be interpreted with caution, and similar patterns have been linked in prior literature to concerns about publication integrity challenges (30–32). We used here also retraction data to penalize the citation impact of institutions against the volume of authors with retracted papers where the retraction has not been due to publisher/journal error. We also used percentile-adjusted information on rates of self-citations and publications in discontinued titles to further calibrate the profile of each institution. Therefore, the generated summary score considers both impact and aspects of research integrity.

Some institutions may have high concentrations of top-cited authors while also showing signals that warrant closer integrity-related examination. Other recent analyses by *Nature* (33) have also revealed that some institutions, especially in China, Saudi Arabia, India, Pakistan, and Ethiopia are indeed retraction hotspots. Accounting for retractions, and inordinate self-citations and publications in discontinued titles can substantially alter the placement of some institutions in the adjusted rankings. We hope that inspection of our data may thus strengthen proper incentives, weaken the drive for publication and citation misconduct, and identify institutions and author groups that require more in-depth scrutiny. Different countries may have macro-environments that generate different types of problems. For example, inspection of the list of institutions with the highest penalties shows that many Chinese institutions are penalized for high retractions, while many Russian institutions are penalized for high self-citations and high percentage of publications in discontinued titles. Overall, in our baseline analysis, Saudi Arabia, China, Malaysia, Iran, India and Indonesia have cumulative penalties that on average exceed their number of top-cited authors. However, as already discussed, none of the three proxies offer full proof of research integrity problems at the institution level, let alone the country level. While they can serve as warning signals, alternative explanations may also exist why their values are high in specific countries, including field mix, coverage bias, diverse affiliation structures, retraction detection differences, and bonus-affiliation practices.

There are several limitations in our work. First, while the precision and recall of Scopus are high, some authors may be misclassified regarding top-cited status because of inaccuracies. Second, assigning a single affiliation to each author also carries uncertainty and the potential for error. Institutions interested in more in-depth assessments of their people may thus wish to verify data against their up-to-date personnel files. For small institutions, we caution that even small errors can cause substantial changes in the number and proportion of highly cited authors. Therefore, while we make available the full data on 6,979 institutions, only the largest one-third of them may be immune to the impact of even very modest errors. Uncertainty for each of the studied proportions can be estimated by standard binomial proportion 95% confidence intervals. Third, we have used an aggregation system for organizations and there can be disagreements on specific aggregations. Requests for re-evaluation and revision of aggregation decisions should be sent directly to Scopus, as we do not personally have permission to make any changes in the Scopus system ourselves. Notably, some institutions currently are under pressure to merge (34,35). While there may be some sensible reasons for such mergers, e.g. reduction of redundancy in resources or cost, other reasons include misaligned incentives – in particular gaining higher ranks in university rankings that depend on volume and size. Fourth, retracted papers are probably only a small fraction of the fraudulent and highly flawed papers that should be retracted (36). We cannot exclude that the fraction is different across different institutions. However, it is impossible to estimate that fraction. Therefore, use of the observed retractions is the best available option. Fifth, some retractions may be due to honest error (37); and scientists recognizing their errors and correcting the record should be applauded. However, retraction notices are usually too elliptical to separate securely honest error from misconduct, and the latter is probably the large majority (38). Finally, while we provide theoretical justification for our adjustment and correction approach to generate a summary score, eventually the exact weighting of the adjustments and corrections is a largely arbitrary choice. Within a reasonable sensitivity range, different corrections for self-citations, discontinued titles, and retractions do not materially alter the overall institutional ordering, but a subset of institutions can still change substantially. Moreover, if science-wide data becomes available for other proxies of research integrity, these may also be considered in the future for additional multi-dimensional institutional assessments.

Overall, the data that we make publicly available may help improve transparency for institution-level impact and research integrity issues among the authors who are currently affiliated with each institution. Careful balance of excellence and research integrity is needed.

## SUPPLEMENTARY TABLES AND FIGURES

All supplementary tables and figures have been posted at Data Mendeley at doi: 10.17632/jn5j7gjpzj.1

## SUPPLEMENTARY TEXT

### APPENDIX METHODS

#### Aggregation of affiliations

Scopus displays a single organization to each Scopus author ID. For authors who have multiple organizations in their published corpus, only one organization is selected based on an automated algorithmic procedure that prioritizes the last listed organization in the most recent papers. Authors can also curate their data and select their own preferred organization based on the list of recent organizations. Authors belonging to the same institution may nevertheless use different ways to present their organization in their work. This may result in heterogeneity and dispersal of the authors of the same institution across different organizations. For example, a scientist affiliated with Harvard, may be shown with their organization being the university (“Harvard University”, n=3699 authors), the school of the university (e.g. “Harvard Medical School” n=15,016, or “Harvard T.H. Chan School of Public Health” n=1,872, or “Harvard Business School” n=283) or occasionally even more circumscribed entities and units within the university (e.g., “The Rowland Institute at Harvard” n=47, or “Smithsonian Astrophysical Laboratory” n=68).

For this project, we use these granular organizations, as reported in the Scopus Author ID for each author to generate “aggregated” organizations which we refer to as institutions. For the institutions all underlying (child) organizations that pertain to the same institution are combined, with the exception that organizations that name only hospitals are kept separate. For example, the institution for Harvard University aggregates 50 different organizations that have at least one author assigned to them among the 10,933,183 authors. However, it does not aggregate hospitals that may have Harvard affiliation. Organizations such as “Massachusetts General Hospital”, “Massachusetts General Hospital Cancer Center”, and “MGH Institute of Health Professions” are merged under a separate institution named Massachusetts General Hospital. A small number of organizations may be assigned to multiple different institutions; these are referred to as “jointly owned” organizations.

The full hierarchy of organizations that are mapped to an institution upon aggregation can be found in the subscription edition of Scopus on the organization profile page by navigating to the “structure” tab. The inclusion and exclusion criteria of these organizations are based on the relationship between the parent organizations and their child-organizations in the hierarchy. Relationships between organizations can be attributable or non-attributable. In the case of an attributable relationship, all output of the child-organization is rolled up to the parent organization and the child-organization is visible in the institutional hierarchy on Scopus. This tree-view of an organizational hierarchy can contain multiple levels of organizations, in other words child-organizations may also have their own child-organizations.

Organizations are more granular than the institutions authors are linked to as these largely reflect the choice of how each author represents themselves (which organization they have listed on their publications). The institutional aggregation achieves far better standardization in this regard, but is also not perfect, e.g. some authors with “Massachusetts General Hospital” listed organizations may have appointments at Harvard University, like those who get aggregated under “Harvard University”, and vice versa.

### APPENDIX RESULTS

#### Single recent-year analyses

Analyses using the single recent-year instead of career-long data provided qualitatively similar inferences (see Supplementary Tables 6-10). Numbers and proportions of top-cited authors were on average slightly higher, given that the list of top-cited authors includes more authors in the recent-year list than in the career-long list. The increase was disproportionately higher for some institutions, especially some of the largest and most high-impact ones (e.g. Harvard and Stanford), and furthermore most prominent for Chinese universities. Focusing on single recent-year impact propels several small institutions from non-high-income countries to the top ranks of proportion of top-cited authors (Supplementary Table 8). However, almost all these institutions are markedly affected by authors with retracted papers and high self-citation and discontinuation rates suggesting gaming of metrics and misconduct in their environment. Focusing on institutions with at least 100 eligible authors for the primary analysis (Supplementary Table 9) removes the very small spurious institutions, but several mid-size institutions predominantly from Saudi Arabia and some from Egypt, India, and United Arab Emirates appear at the very top ranks in the absence of adjustments, but almost all of them lose their top-ranks or even end up with negative adjusted numbers. This may reflect gaming practices (as captured by retractions and high self-citations) or the common use of Saudi Arabian institutional affiliations by scientists in other countries who are paid heavy bonuses to do this.

#### Sensitivity analyses

Sensitivity analyses excluding from the primary analysis authors who had not actively published since 2020 removed about 12% of authors from the analysis but yielded qualitatively similar inferences (not shown).

Across all sensitivity analyses on different p and f adjustments, institutional percentile rankings remained highly correlated with the baseline specification (Supplementary Figures S1-8). As shown in Supplementary Figure S1 for the career-long impact, Spearman correlations ranged from 0.928 to 0.991 across all combinations of percentile cutoffs (p = 80, 95, 99) and retraction-penalty factors (f = 1, 2, 4) with the baseline of p=95 and f=2. Bland-Altman analyses (Supplementary Figure S2) confirm that most institutions deviate only modestly from baseline, although a small minority demonstrate shifts of approximately 20–40 percentile points under the most extreme correction settings. Scatterplots of adjusted percentages (Supplementary Figure S3) show equally strong agreement, and KDE plots (Supplementary Figure S4) demonstrate that the overall distribution of adjusted institutional scores changes only minimally under alternative parameter settings.

Sensitivity analyses for the single-year citation impact produced qualitatively similar patterns (Supplementary Figures S5-8) but with slightly lower correlations and somewhat greater dispersion (Spearman correlations ranging from 0.897 to 0.986). These sensitivity analyses support the use of the baseline specification (p = 95, f = 2) as a pragmatic reference choice, as results remain largely consistent across a wide range of alternative parameter settings.

## REFERENCES

1. Ioannidis JP, Patsopoulos NA, Kavvoura FK, Tatsioni A, Evangelou E, Kouri I, Contopoulos-Ioannidis DG, Liberopoulos G. International ranking systems for universities and institutions: a critical appraisal. BMC Med. 2007;5:30.

2. Vernon MM, Balas EA, Momani S. Are university rankings useful to improve research? A systematic review. PLoS ONE 2018;13(3):e0193762.

3. Billaut J-C, Bouyssou D, Vinckecoara P, Should you believe in the Shanghai ranking? An MCDM view. Scientometrics 2010;84:237–63.

4. Coalition for Advancing Research Assessment, in: https://www.coara.org/, last accessed November 21, 2025.

5. Waltman L, Calero-Medina C, Kosten J, Noyons ECM, Tijssen RJW, Jan van Eck N, et al. The Leiden ranking 2011/2012: Data collection, indicators, and interpretation. Journal of the American Society for Information Science and Technology 2012;63:2419–2432.

6. Baas J, Schotten M, Plume A, Cote G, Karimi R. Scopus as a curated, high-quality bibliometric data source for academic research in quantitative science studies Open Access. Quantitative Science Studies 2020;1(1):377–386.

7. Ioannidis JPA, Baas J, Klavans R, Boyack KW. A standardized citation metrics author database annotated for scientific field. PLoS Biol. 2019;17(8):e3000384.

8. Ioannidis JP, Klavans R, Boyack KW. Multiple Citation Indicators and Their Composite across Scientific Disciplines. PLoS Biol. 2016;14(7):e1002501.

9. Ioannidis JPA, Boyack KW, Collins TA, Baas J. Gender imbalances among top-cited scientists across scientific disciplines over time through the analysis of nearly 5.8 million authors. PLoS Biol. 2023;21(11):e3002385.

10. Ioannidis JPA, Boyack KW, Baas J. Updated science-wide author databases of standardized citation indicators. PLoS Biol. 2020;18(10):e3000918.

11. August 2025 data-update for “Updated science-wide author databases of standardized citation indicators”, in: https://elsevier.digitalcommonsdata.com/datasets/btchxktzyw/8, last accessed November 21, 2021.

12. Kawashima H, Tomizawa H. Accuracy evaluation of Scopus Author ID based on the largest funding database in Japan. Scientometrics 2015;103:1061–71.

13. Archambault E, Beauchesne OH, Caruso J. “Towards a multilingual, comprehensive and open scientific journal ontology” in Proceedings of the 13th International Conference of the International Society for Scientometrics and Informetrics (ISSI), Durban, South Africa. Noyons B, Ngulube P, Leta J, editors. 2011: 66–77.

14. Ioannidis JPA. Features and signals in precocious citation impact: A meta-research study. PLoS One. 2025;20(8):e0328531.

15. Meho LI. Gaming the metrics: bibliometric anomalies in global university rankings and the research integrity risk index (RI2). Scientometrics 2025;130:6683–6726.

16. Retraction Watch Database, in: http://retractiondatabase.org/RetractionSearch.aspx?, last accessed November 21, 2025.

17. Ioannidis JPA, Pezzullo AM, Cristiano A, Boccia S, Baas J. Linking citation and retraction data reveals the demographics of scientific retractions among highly cited authors. PLoS Biol. 2025;23(1):e3002999.

18. Boccia S, Cristiano A, Pezzullo AM, Baas J, Roberge G, Ioannidis JP. Gender imbalances of retraction prevalence among highly cited authors and among all authors. bioRxiv 2026, doi: 10.1101/2025.07.08.662297

19. Van Noorden R, Singh Chawla D. Hundreds of extreme self-citing scientists revealed in new database. Nature 2019;572(7771):578–9.

20. Holland K, Brimblecombe P, Meester W, Chen T. The importance of high-quality content and reevaluation in Scopus. In: https://assets.ctfassets.net/o78em1y1w4i4/4e0v9wh0vDmzyNnXTh0cCf/31eac680074c579a5c33fda620efa093/The-importance-of-high-quality-content-curation-and-reevaluation-in-Scopus.pdf, 2021. Last accessed April 20, 2026.

21. Jin GZ, Jones B, Lu SF, Uzzi B. The reverse Matthew effect: consequences of retraction in scientific teams. Rev Econ Stat. 2019;101(3):492–506.

22. Azoulay P, Furman JL, Krieger JL, Murray FE. Retractions. Rev Econ Stat. 2014;96 doi: 10.1162/REST_a_00469.

23. Gadd E. Mis-measuring our universities: why global university rankings don’t add up. Front Res Metr Anal. 2021;6:680023.

24. Soh K. The seven deadly sins of world university ranking: a summary from several papers. Journal of Higher Education Policy and Management 2017;39:104–115.

25. Saisana M, d’Hombres B, Saltelli A. Rickety numbers: Volatility of university rankings and policy implications. Res Policy 2011;40:165–77.

26. Hicks D, Wouters P, Waltman L, de Rijcke S, Rafols I. Bibliometrics: The Leiden Manifesto for research metrics. Nature 2015;520:429–431.

27. Abramo D. The forced battle between peer-review and scientometric research assessment: Why the CoARA initiative is unsound. Research Evaluation 2024;

28. Ioannidis JPA, Maniadis Z. In defense of quantitative metrics in researcher assessments. PLoS Biol. 2023;21(12):e3002408.

29. Ioannidis JPA, Maniadis Z. Quantitative research assessment: using metrics against gamed metrics. Intern Emerg Med. 2024;19(1):39–47.

30. Ro C, Leeming J. Authorship for sale: Nature investigates how paper mills work. Nature 2025;642(8068):823–826.

31. Sebo P. Chinese authors are overrepresented in medical articles retracted for fake peer review or paper mill. Intern Emerg Med. 2024 Nov;19(8):2369–2371.

32. Sebo P, Sebo M. Geographical Disparities in Research Misconduct: Analyzing Retraction Patterns by Country. J Med Internet Res. 2025 Jan 14;27:e65775.

33. Van Noorden R. These universities have the most retracted scientific articles. Nature 2025;638(8051):596–599.

34. Aula HM, Tienari J. Becoming “world class”? Reputation building in a university merger. Critical Perspectives on International Business 2011;7:7–29.

35. Docampo D, Egret D, Cram L. The effect of university mergers on the Shanghai ranking. Scientometrics 2015;104:175–191.

36. Oransky I. Retractions are increasing, but not enough. Nature 2022;608(7921):9.

37. Wang D, Chen S. An empirical study of retractions due to honest errors: Exploring the relationship between error types and author teams. J Informetrics 2025;19:101600.

38. Hwang SY, Yon DK, Lee SW, Kim MS, Kim JY, Smith L, Koyanagi A, Solmi M, Carvalho AF, Kim E, Shin JI, Ioannidis JPA. Causes for retraction in the biomedical literature: a systematic review of studies of retraction notices. J Korean Med Sci. 2023;38(41):e333.

